# Surface electromyography (sEMG) of equine core muscles and kinematics of lumbo-sacral joint during core strengthening exercises

**DOI:** 10.1101/2023.08.11.552791

**Authors:** Judit Aulinas Coll, Scott Blake, Roberta Ferro de Godoy

## Abstract

Dynamic Mobilisation Exercises (DME) and myotatic reflex exercises were developed with the aim of improving core strengthening in horses. Previous studies have shown DME can increase cross sectional area (CSA) and symmetry of multifidus muscle, as well as activating the external oblique abdominal, and superficial descending pectoral muscles. The aim of this study was to objectively measure activity differences in *m. longissimus dorsi* (LD) and *m. rectus abdominus* (RA) whilst performing three levels of spinal flexion and lateral bending, as well as comparing thoracic and pelvic lift exercises in nine adult sport horses. Three repetitions of each exercise was performed for five seconds. Surface electromyography (sEMG) was used to record muscle electric activity, whilst sagittal lumbo-sacral flexion was measured with kinematics analysis. Overall, the results have shown that spinal flexion and lateral bending activate the *m. rectus abdominis* (RA) progressively as the exercise requires further reach, with a lateral bending effect evident on the ipsilateral side of RA. RA also had increased activation during thoracic lifts in comparison with pelvic lifts. *M. longissimus dorsi* (LD) has shown no significant differences in peak or average rectified EMG measures on the contralateral side during lateral bending. Pelvic lifts generated the greatest flexion of the lumbo-sacral (LS) joint. Results provide a guideline of the level of muscle effort required in relation to each exercise.

**Highlights:**

- *M. rectus abdominis* is activated more with further reach for cervical flexion and lateral bending.
- *M. rectus abdominis* is more active during thoracic lift than pelvic lift.
- Lumbo-sacral joint achieves greater flexion on pelvic lift exercises.
- *M. longissimus dorsi* responds very little to increased reach in DMEs.

## 1. Introduction

Core training should be performed throughout a horse’s athletic career to maintain a healthy back, or used as a therapeutic intervention when back pain is identified [1]. Dynamic mobilisation exercises (DME) are core strengthening exercises for horses designed by Stubbs and Clayton [2], which differ from passive stretches as they require concentric contraction as well as isometric contraction to maintain balance[1]. DMEs include rounding and lateral bending exercises, which are both commonly used for core muscle activation[3].Myotatic reflexes, as belly and pelvic lifts, are another category of core strengthening exercises. These exercises are designed to target postural muscles to promote balance and self-carriage. Belly and pelvic lifts rely on the horse’s response to pressure applied to specific anatomic areas, by bending of specific intervertebral joints through activation of the long mobilizing muscles[1]. According to Stubbs and Clayton [2], thoracic/belly lifts leads to flexion on the thoracic area, whilst pelvic lift/tuck flexes and lifts the lumbar and lumbosacral (LS) joints. It is expected that the higher range of motion on sagittal motion on the lumbar area joint occurs at LS joint due to the thickness and decreased height of its intervertebral disc, as well as the absence of real supraspinous ligament [4]Therefore, these exercises could help to maintain or regain LS flexion/extension range of motion, which is important for performance, especially during running gaits [5]

Therapeutic use of DMEs have been shown to increase the cross-sectional area (CSA) and restore symmetry of the *m. multifidus spinae*[6–9], increase the CSA of *m. longissimus dorsi* [6] and enhance overall posture [10]. *M. multifidus* atrophy has been associated with back pathologies [11] and chronic limb lameness[12], and typically does not recover spontaneously. Persistent atrophy of the deep stabilizing muscles seems to be common[13], indicating a need for core training exercises to regain symmetry. DMEs can be used to correct these deficits by addressing the muscular and neurological system, assisting with re-education of proprioception and addressing activation of the deep back stabilising muscles. Aside from the benefits on rehabilitation of axial and limb injuries [13], core exercises have also been shown to improve recovery and an earlier return to work in horses post colic surgery [14].

Evidence-based approaches and objective outcome measurements in equine rehabilitation are of major importance, so the use of non-invasive surface electromyography (sEMG) technology to assess muscle function is an important tool. sEMG has been used to assess the different therapeutic exercises effects on core muscles, such as pole work, elastic resistance bands [15], and Pessoa training aids [16], however, research regarding core muscle activation with DMEs is still scarce. Destabilisation exercises have shown to increase sEMG signals in the *m. pectoralis descendens* whilst sustained chin-to-hip lateral bending and chin-between-knees flexion has been shown to increase activity on *m. obliquus externus abdominis* [3]. Exercise using thoracic and pelvic lifts, as well as combing with a tail pull, on *m. longissimus dorsi, m. rectus abdominis and m. gluteus medius* can elicit an increase in *m. rectus abdominis* activation and *m. longissimus dorsi* activity [17]. However, to the authors’ knowledge, no studies have assessed the effect of different DMEs in the *m. rectus abdominis and m. longissimus* dorsi activity simultaneously, and at the different levels of exercise. The aims of this research were therefore to assesses *m. longissimus dorsi* and *m. rectus abdominis* peak and average rectified EMG measures on three levels of spinal flexion and lateral bending exercises and two types of myotatic reflexes: thoracic and pelvic lifts, to determine which exercise and/or level would lead to the higher activation of these muscles. In addition, we also wished to determine which exercise would cause higher flexion within the lumbo-sacral joint, as its health is also essential to performance.

## 2. Material & Methods

The material in this manuscript has been acquired according to guidelines set by The Animal (Scientific Procedures) Act 1986 and has been approved by the Animal Welfare and Ethics Committee of Writtle University College. The approval number is 1300/2021. A written informed consent was obtained from the owners of the participants of the study.

### 2.1. Animals

The studied population consisted of a convenience sample of nine horses (n=9), four mares and six geldings (age: 12.3 ± 4.94 years old; height 161.44 ±5.45cm), including Irish Sport Horses, Warmbloods and Thoroughbreds. All horses were training and/or competing in dressage or show jumping. The selection criteria excluded horses with previous history of back pain/pathologies or with signs of pain and lameness, young horses, horses not trained or unable to perform the DMEs. Prior to data collection, all horses underwent a training protocol consisting of five days performing dynamic mobilisation exercises to ensure correct performance. The training was followed by two days rest to recover from muscle fatigue.

### 2.2. Core strengthening exercises Protocol

Different core strengthening exercises, based on Stubbs and Clayton [2], were considered for this protocol (Figure 1). In all exercises, horses were standing in a square position on a flat, non-slip surface. Exercises included three spinal flexion levels: chin-to-chest, chin-to-carpus and chin-to-fetlock; three spinal lateral bending levels, bilaterally: chin-to-shoulder, chin-to-girth and chin-to-hip; and two myotatic reflex back lift exercises: thoracic and pelvic lifts.

**Figure 1.** Representation of the different dynamic mobilisation exercises performed. Top row: spinal flexion exercises, from left to right: chin-to-chest, chin-to-carpus, and chin-to-fetlock. Middle row: lateral bending exercises, from left to right: chin-to-shoulder, chin-to-girth, and chin-to-hip. Bottom row: myotatic reflex exercises, from left to right: belly lift, and pelvic lift.

Each horse was encouraged to perform each exercise using a bait, molasses lick tub or carrot, depending on preference. For the spinal flexion, the horse was encouraged with the bait to place the chin downwards and pass the head between the chest, knees and fetlock, with the latter considered to be the highest reach of exercise. During lateral bending, horses were encouraged to bend the neck and head laterally to the shoulder, girth and hip, deemed as being from a lower to higher level of reach. Each exercise was performed to the left and right side for the lateral exercises, whilst each mobilisation was repeated three times during an exercise session, with each position held for five seconds.

Lift exercises are considered to be those that recruit and strengthen the abdominal and pelvic-stabilizing muscles in response to a tactile pressure[8]. Thoracic lift was encouraged by sliding two hands with a moderate amount of pressure caudally over the sternum line. Pelvic lift was performed by manually applying pressure to a point located between *m. biceps femoris* and *m. semitendinosus*. Horses were required to hold each lift for five seconds, with three repetitions per session. An interval of one minute between repetitions was given to avoid any muscle warm up effect or fatigue.

Figure 1 shows the representation of all exercises performed.

### 2.3. Surface Electromyography (sEMG)

A sEMG (Neurotrac MyoPlus 2 Pro, Verity Medical Ltd, Hampshire, UK), with 0 to 2000 μV RMS continuous range, and sensitivity of 0.1 μV RMS, was used alongside its dedicated computer software for analysis of the left *m. rectus abdominis* (RA) and left lumbar *m. longissimus dorsi* (LD) activity. Preparation included clipping the lumbar region of the left LD (between L2 and L4) and caudal to sternum for the left RA with both clippers and a razor. Skin patches were cleaned and wiped with surgical spirit to remove any grease and dirt to enable a better connection to the electrode pads[16]. The same researcher (JAC) clipped the skin to ensure that electrode placement was correct and consistent across all horses. A pair of pre-gelled electrodes (50 x 50mm square pads) were placed 5 cm lateral from the spine at the left lumbar level (one electrode between L2-L3 and another between L3-L4). Another pair of electrodes were placed 2 cm from the *linea alba*, caudal to the sternum, on the left RA belly. (Figure 2) The space inter-electrode was 1cm, which is within the range of what is utilized in other equine sEMG studies (range 1 to 4cm) [15,18]. A reference electrode was also placed over the left *tuber coxae*, [19]. The order of each exercise was randomized, and each exercise was repeated three times. Data was analysed using manufacturers software (Neurotrac software, Verity Medical Ltd, Hampshire, UK). Peak and average muscle activity in μV for each condition were recorded. For the lateral bending exercises, the muscle activity was measured when the assessed muscle was on the ipsilateral (lateral bending to the left) and the contralateral side (lateral bending to the right) of the bending. Figure 3 shows a representation of sEMG data collected during three repetitions of chin-to-fetlock exercise.

**Figure 2.**
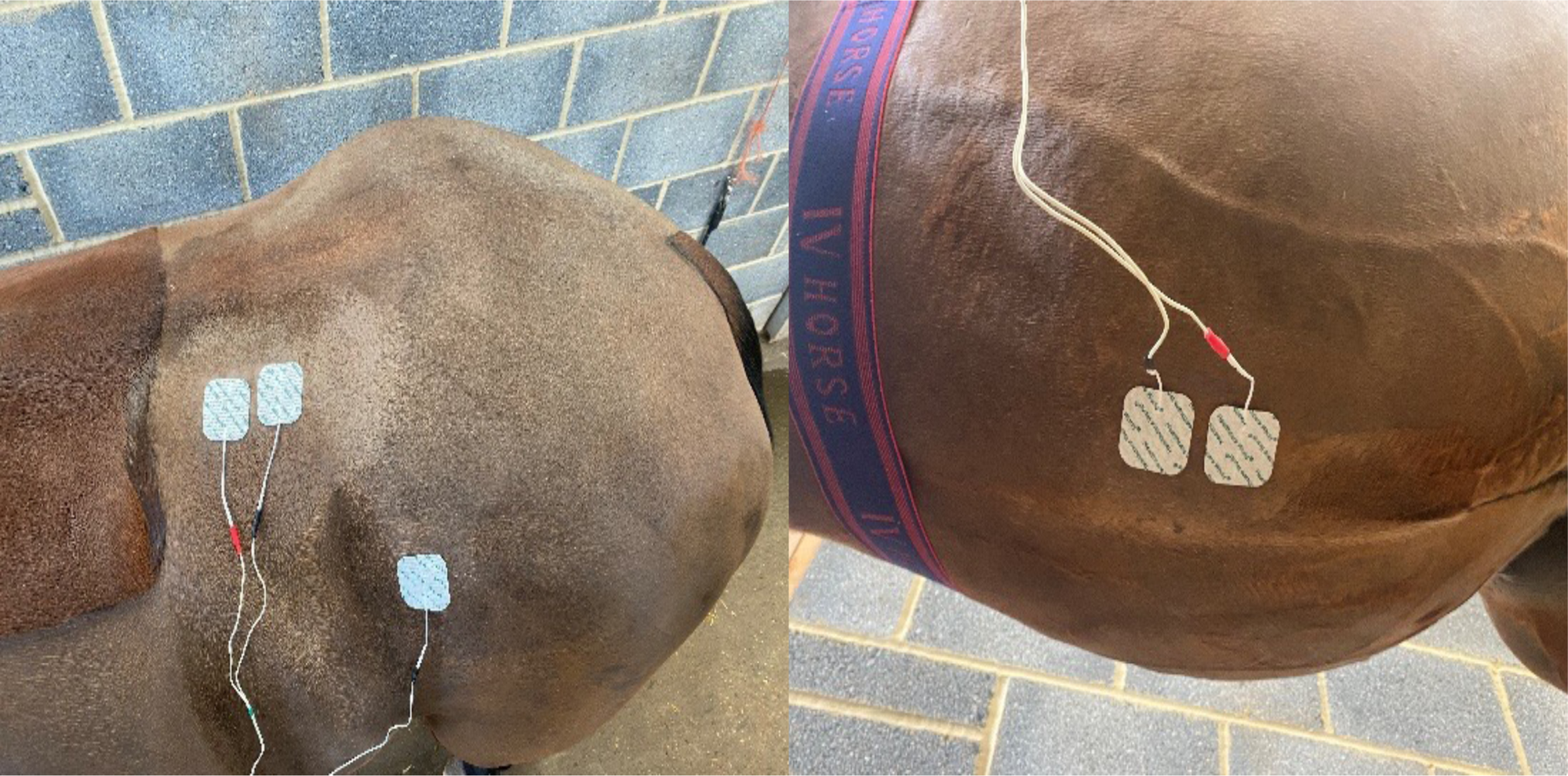
Positioning of sEMG electrodes on the *m. longissimus dorsi* (left) and *m. rectus abdominis* (right).

**Figure 3.**
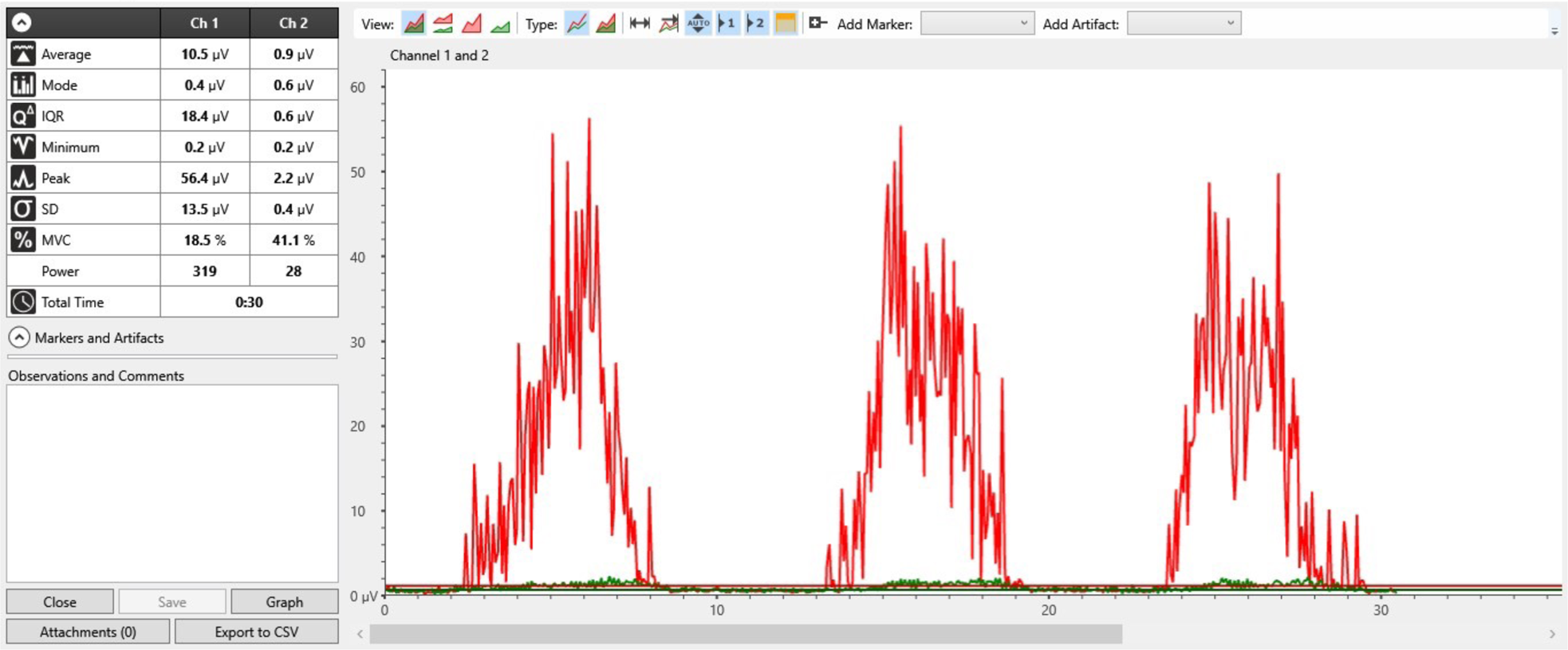
Plot of data acquired from sEMG. The graph shows the muscle activity of a horse performing three repetitions of chin-to-fetlock spinal flexion dynamic mobilisation exercise. Red graph represents *m. recuts abdominus*, green graph represents *m. longissimus dorsi*.

Raw sEMG signals were differentially amplified by a common mode rejection ratio of 130 dB/s and a bandpass filter of 3dB bandwidth wide (18 ± 4 Hz cut-off) and narrow filter (370 ± 10 Hz cut-off) and a notch filter of 50Hz. sEMG variables included average rectified value (ARV) and peak sEMG measures for each muscle across all exercises. ARV and peak sEMG measures were calculated by the dedicated software using rectified signals with exercise duration as temporal domain. Outliers in ARV data were detected and removed by setting upper and lower outlier limits as two standard deviations outside of the mean ARV values within each horse, muscle, and condition according to described by St. George et al. [20] Outliers in peak amplitude data were detected from baseline conditions prior to normalization of continuous signals, using the same method employed for ARV data, and removed.

### 2.4. Lumbo-sacral flexion-extension kinematics

Kinematics data was collected whilst each horse was on a flat, even surface. Reflective markers 3cm diameter were placed on the spinous process of the fifteenth thoracic vertebrae, lumbo-sacral joint and great trochanter of femur, as per Walker et al. [21]. Horses were recorded perpendicular to a single high-speed video camera mounted in a level position on a tripod, approximately 3 m from horse, with one LED spotlight (500 W) to illuminate the reflective markers. The camera collected data at 240 Hz (resolution 1,334×750 pixels, 720p HD), with a field of view capturing the entire horse whilst performing all exercises. The videoing started with the horse being placed standing square, with the head and neck position maintained in a neutral posture and the horse’s mouth held level with the point of the shoulder by an assistant. This was considered the lumbo-sacral joint baseline angle. The exercises were then performed whilst the lumbo-sacral kinematics response was being recorded.

Motion analysis software (Quintic Biomechanics v.33, Quintic Consultancy Ltd. Birmingham, UK) was used to analyse each video. The maximum difference between the lumbo-sacral angle (in degrees) in neutral position vs. performing the exercise was calculated. A negative number demonstrated an extension of the lumbo-sacral joint whilst a positive number denotes flexion on the lumbo-sacral joint in relation to the neutral position. Each exercise was repeated three times with the average of the three angles was used for statistical analysis.

### 2.5. Statistical Analysis

Once all outcome variables were analysed, the data was tabulated in excel, and subsequently transferred to SPSS (v28.0, IBM).

The EMG data and lumbo-sacral kinematics for each exercise were subjected to a Shapiro-Wilk test for normality, which showed mostly non-parametric data. All statistical tests comparing cervical flexion levels or lateral bending levels were performed with Friedman test to test differences between the three levels of DMEs A post hoc analysis was conducted when a significant difference was encountered, with Bonferroni correction (95% confidence interval (CI)) for multiple pairwise comparisons. The results report SPSS Bonferroni adjusted p-values. To compare ipsilateral and contralateral muscle activations within the lateral bending exercises, Wilcoxon’s rank test was used. And finally, the pelvic and thoracic tilts data has shown to be of parametric distribution, therefore a paired t-test was used to compare these exercises. Significance level was set at 95% (p<0.05).

## 3. Results

### 3.1. M. rectus abdominis

On the cervical flexion exercises, there were differences in RA ARV sEMG measures (X^2^(2)=8.471, p=0.014) between the DME levels. RA ARV sEMG was significantly higher on chin-to-fetlock exercise than chin-to-chest (p=0.014) (Figure 4). The same trend was seen for peak sEMG measure of RA during cervical flexion DMEs (X^2^(2)=12.667, p=0.002). With the chin-to-fetlock DME having a higher peak sEMG signal than chin-to-chest (p=0.001) (Figure 4).

**Figure 4.**
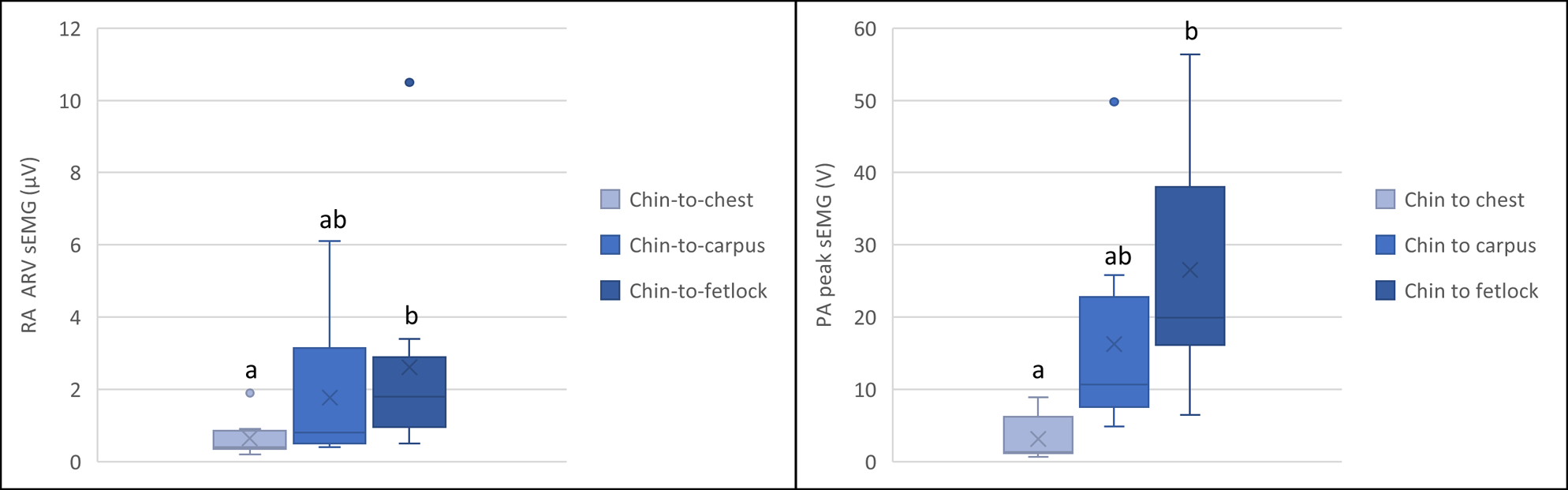
Box plots for ARV (left) and peak (right) *m. rectus abdominus* sEMG measures during different spinal flexion exercises. The bottom and top of the box are the first and third quartiles, and the band inside the box is the second quartile (the median). The cross is the mean. The lines extending vertically from the boxes (whiskers) indicate the minimum and maximum of all of the data. Letters represent significant differences between exercise intensities (p<0.05) (n=9).

For the lateral bending, the contralateral RA, in relation to the bending side, was not statistically significantly different between the three levels of lateral bending neither for ARV sEMG (X^2^(2)=4.941, p=0.085) nor for the peak sEMG signal (X^2^(2)=4.222, p=0.121) (Figure 5). However, for the ipsilateral RA, the muscle sEMG signal was significantly different for ARV (X^2^(2)=16.222, p<0.0001) and peak (X^2^(2)=9.556, p=0.008) sEMG signals, being higher at chin-to-hip DME in comparison with chin-to-shoulder, for both ARV (p=0.000185) and peak (p=0.007) muscle sEMG measures (Figure 5).

**Figure 5.**
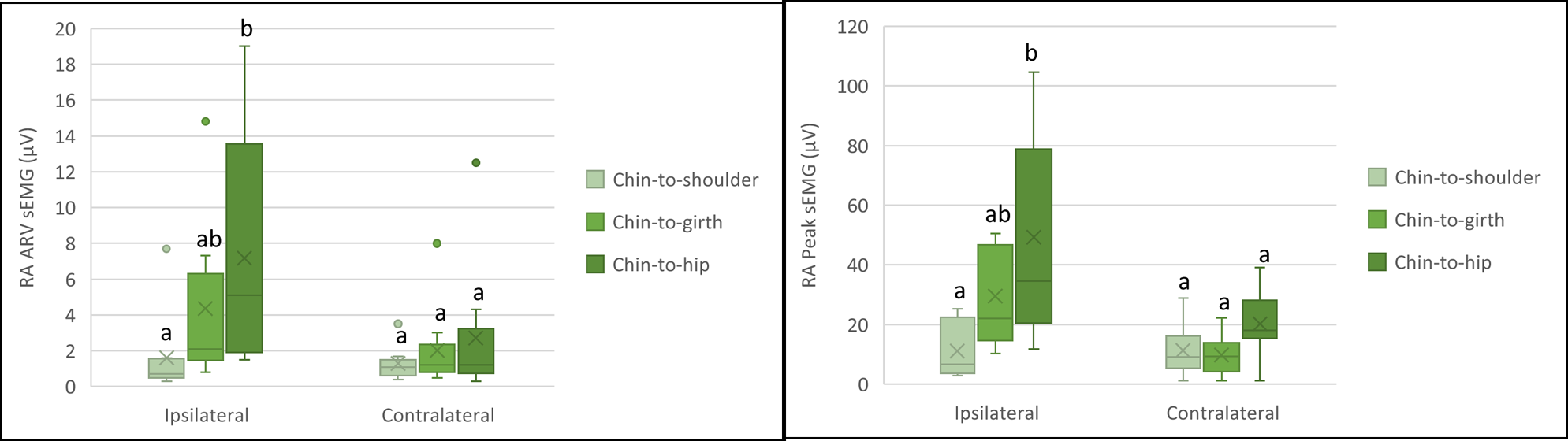
Box plots for ARV (left) and peak (right) *m. rectus abdominus* (ipsilateral and contralateral sides) sEMG measures during different lateral bending exercises. The bottom and top of the box are the first and third quartiles, and the band inside the box is the second quartile (the median). The cross is the mean. The lines extending vertically from the boxes (whiskers) indicate the minimum and maximum of all of the data. Letters represent significant differences between exercise intensities (p<0.05) (n=9).

Thoracic lift also generated a higher peak sEMG measure of RA than pelvic lift (t(7)=2.772, p=0.028) (Figure 6).

**Figure 6.**
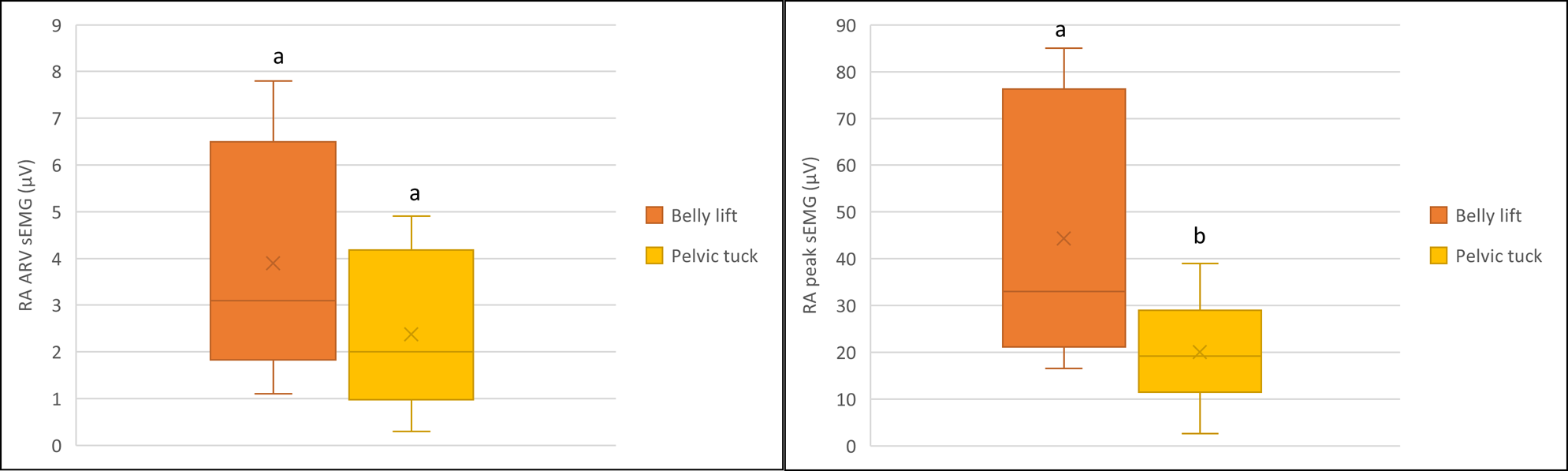
Box plots for ARV (left) and peak (right) *m. rectus abdominus* sEMG measures during belly and pelvic lifts. The bottom and top of the box are the first and third quartiles, and the band inside the box is the second quartile (the median). The cross is the mean. The lines extending vertically from the boxes (whiskers) indicate the minimum and maximum of all of the data. Letters represent significant differences between exercise intensities (p<0.05) (n=9).

### 3.2. M. longissimus dorsi

The LD did not show any differences between levels of the cervical flexion DMEs in ARV (X^2^(2)=5.407, p=0.067) or peak (X^2^(2)=0.667, p=0.717) sEMG signals (Figure 7).

**Figure 7.**
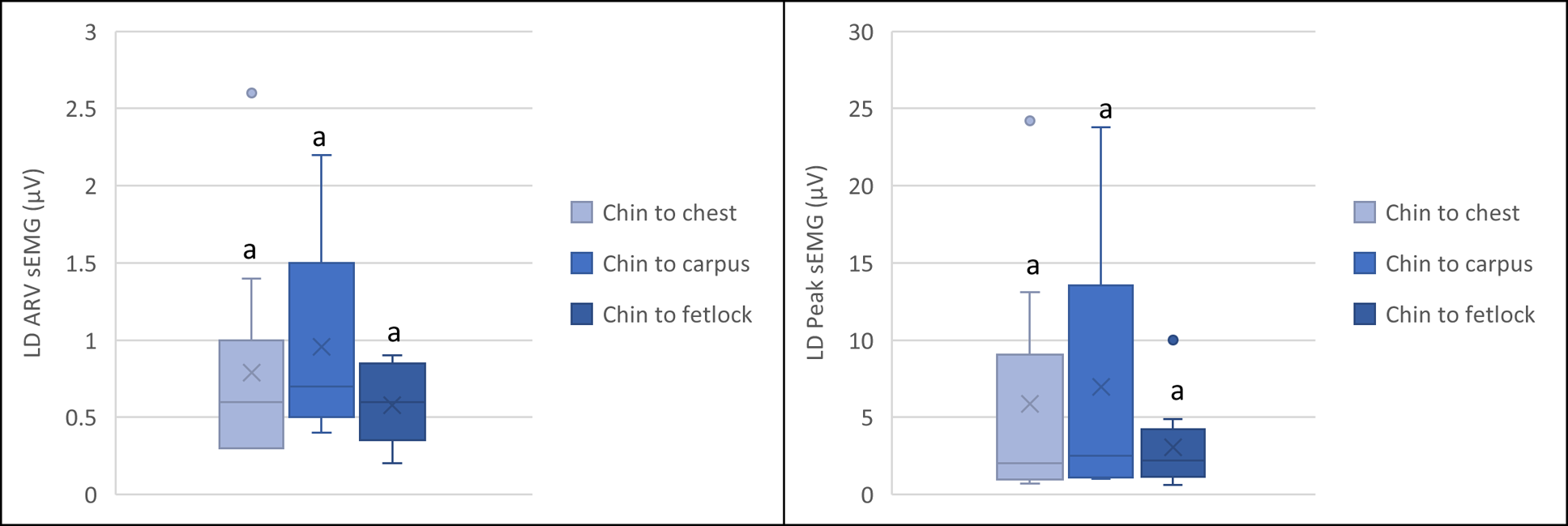
Box plots for ARV (left) and peak (right) *m. longissimus dorsi* sEMG measures during different spinal flexion exercises. The bottom and top of the box are the first and third quartiles, and the band inside the box is the second quartile (the median). The cross is the mean. The lines extending vertically from the boxes (whiskers) indicate the minimum and maximum of all of the data. Letters represent significant differences between exercise intensities (p<0.05) (n=9).

LD peak and ARV sEMG measures on the contralateral LD during different reaches of lateral bending exercises was non-significant (p>0.05) (Figure 8).

**Figure 8.**
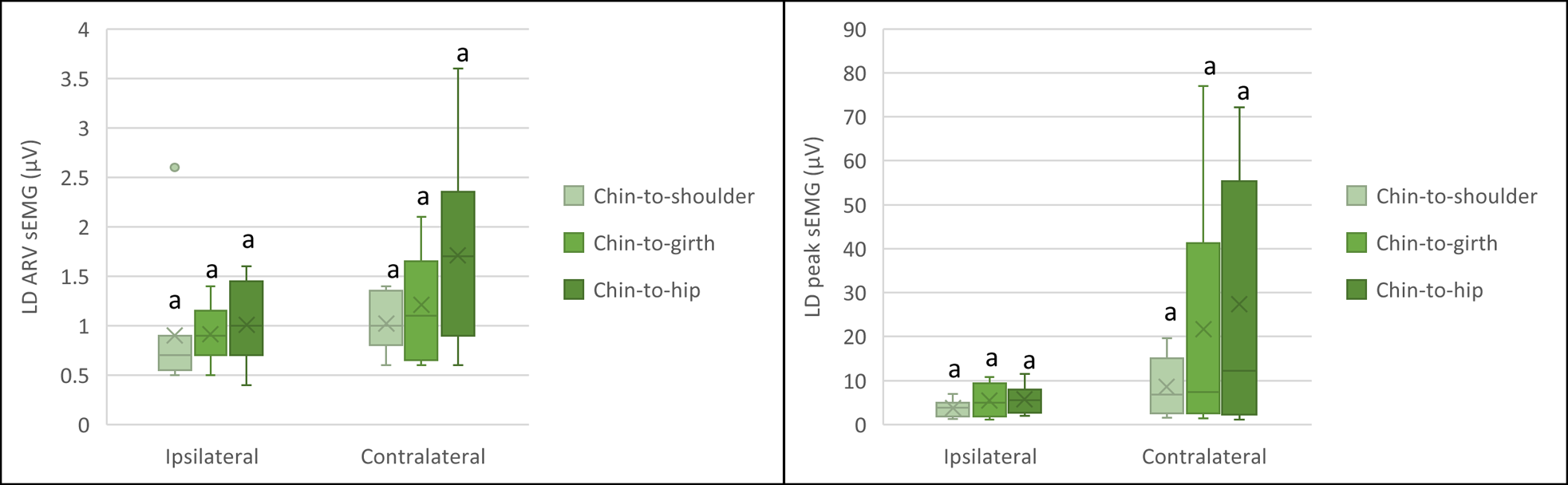
Box plots for ARV (left) and peak (right) *m. longissimus dorsi* (ipsilateral and contralateral sides) sEMG measures during different lateral bending exercises. The bottom and top of the box are the first and third quartiles, and the band inside the box is the second quartile (the median). The cross is the mean. The lines extending vertically from the boxes (whiskers) indicate the minimum and maximum of all of the data. Letters represent significant differences between exercise intensities (p<0.05) (n=9).

LD did not show significant differences in neither peak nor ARV sEMG between thoracic or pelvic lifts (p>0.05) (Figure 9).

**Figure 9.**
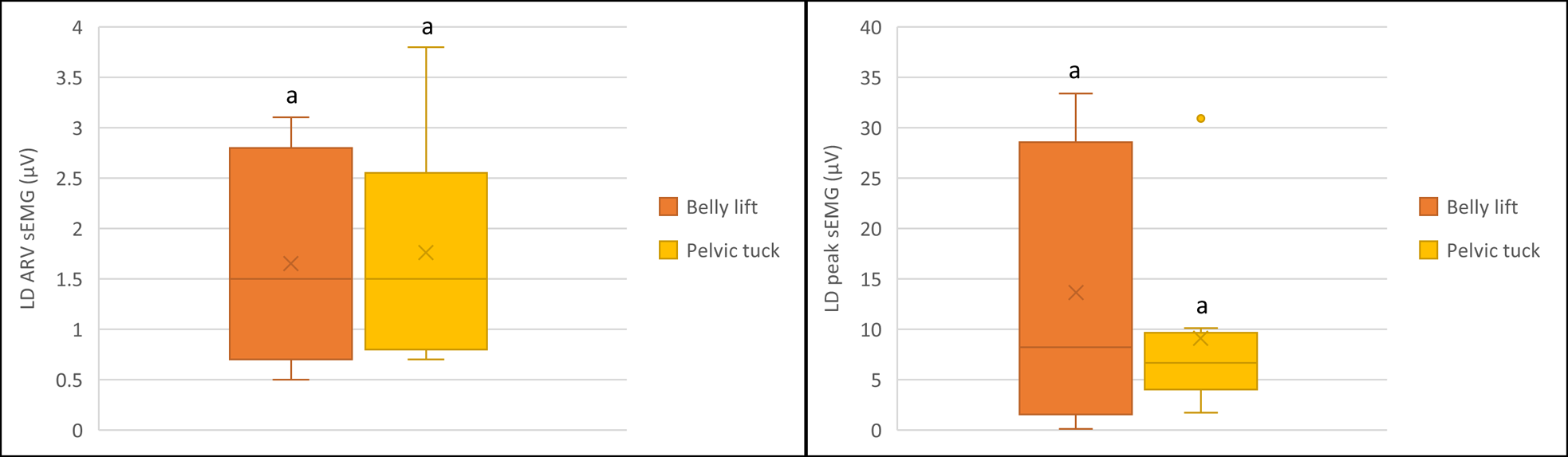
Box plots for ARV (left) and peak (right) *m. longissimus dorsi* sEMG measures during belly and pelvic lifts. The bottom and top of the box are the first and third quartiles, and the band inside the box is the second quartile (the median). The cross is the mean. The lines extending vertically from the boxes (whiskers) indicate the minimum and maximum of all of the data. Letters represent significant differences between exercise intensities (p<0.05) (n=9).

### 3.3. Comparison of RA and LD amongst all core strengthening exercises

Statistical analysis was also performed to compare muscles activation amongst all exercises performed. For ARV sEMG of RA, there was a significant difference between exercises (X^2^(2)=6.645, p=0.036), with chin-to-hip showing the highest average (p=0.037) and peak (p=0.037) sEMG on the ipsilateral RA. For LD, the pelvic tilt was the exercise showing the higher ARV sEMG signal amongst all exercises (p=0.014). No differences were seen between exercises in LD peak sEMG (p>0.05)

### 3.4. Lumbo-sacral kinematics

The lumbo-sacral joint kinematics were analysed to compare how different exercises affect LS joint angles in relation to neutral posture (negative number = extension of LS in relation to the neutral position; positive number=flexion on the LS in relation to the neutral position) (Figure 10).

**Figure 10.**
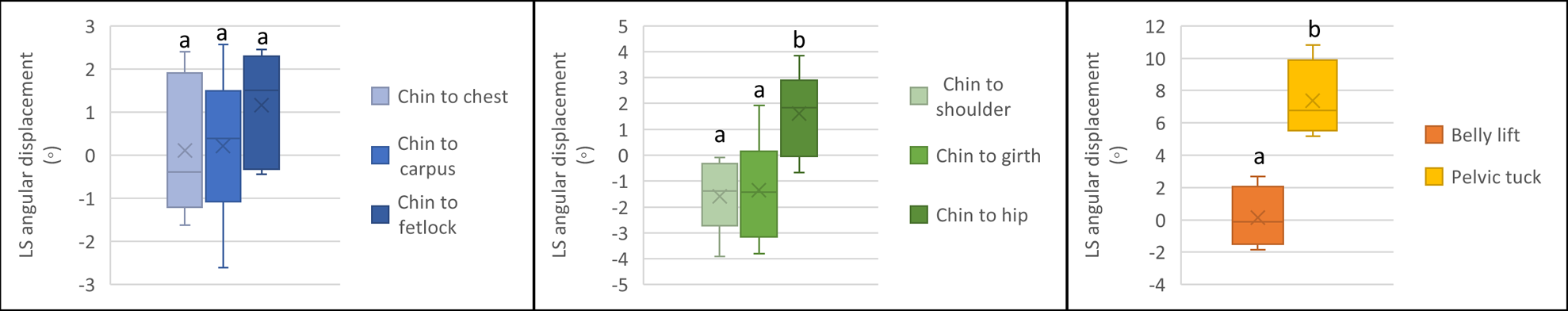
Box plots lumbosacral kinematics during the different exercises performed. The bottom and top of the box are the first and third quartiles, and the band inside the box is the second quartile (the median). The cross is the mean. The lines extending vertically from the boxes (whiskers) indicate the minimum and maximum of all of the data. Letters represent significant differences between exercise intensities (p<0.05) (n=9).

For the flexion exercises, LS joint was more flexed on chin-to-fetlock compared to chin-to-carpus and followed by chin-to-chest. However, these differences were not statistically significant (F(2,10)= 1.680, p=0.235). During lateral bending exercises, LS kinematics was significantly different between the 3 levels of the exercise (F(2,10)=5.752, p=0.022). LS flexion was greater during chin-to-hip exercises, compared to chin-to-girth and chin-to-shoulder (p<0.05). Pelvic lift elicited a significantly higher LS flexion in comparison to thoracic lifts (p=0.031).

## 4. Discussion

This study compared muscle activity (sEMG) and lumbosacral flexion when performing different core strengthening exercises. Overall, the RA showed the greatest response across all exercises, with chin-to-fetlock, chin-to-hip (ipsilateral RA) and thoracic lifts generating the highest peak sEMG within the muscle. The pelvic lift exercise elicited the highest flexion movement on the lumbo-sacral joint. Regarding the LD muscle, none of the exercises showed superior ARV or peak sEMG of this muscle, although the LD had a slightly higher peak sEMG on the contralateral muscle during lateral bending.

Previous studies have classified the DMEs based on changes of cervical angulation [22,23]. Spinal flexion exercises showed progressively increase of angulation from the sixth thoracic vertebrae (T6) to the first lumbar (L1) from chin to chest followed by chin to carpus and chin to fetlock. On spinal bending exercises, chin to hip had a greater angulation from (T6-L1) compared to chin to girth [22,23]

sEMG data processing is complex and the muscle activity can be summarised using different output variables, with the two most common measures being peak EMG and average rectified variable (ARV) sEMG. The peak sEMG variable gives a measure of the maximal activity of the given muscle during the exercise and can be used to quantify muscle activity during core exercises. In contrast, the ARV sEMG is a measure of the area under the normalised sEMG time-series curve divided by the time period, and it is an indication of any submaximal activity which may occur during the stabilisation of the body [24]. Therefore, both methods should be included in any sEMG study on the core musculature and core exercises. In our study, sEMG signal amplitude showed higher peak and ARV sEMG of the RA muscle in all the exercises tested, with chin-to-fetlock generating a greater muscle peak and ARV sEMG measures in relation to the other spinal flexion exercises; chin-to-hip showed the highest RA peak and ARV sEMG amongst the lateral bending exercises; and belly lift created more peak and ARV sEMG of the RA than thoracic lift. This supports the theory that further reach exercises leads to higher muscle activity, meaning exercise reach increase during pre-habilitation and rehabilitation is meaningful. Although previous research had confirmed the activation of abdominal muscles (external abdominal oblique (EAO) [3], to our knowledge, this is the first study to compare the different intensities of spinal flexion and lateral bending exercises. Adding our results to those of Gamucci et al. [3], we can conclude that both RA and EAO are highly active during spinal flexion and lateral bending (ipsilateral muscles for lateral bending).

Surprisingly, the LD muscle showed relatively low ARV during all the exercises and no significant differences were found between exercises. An unexpected finding was the increase signal amplitude seen on the contralateral LD during lateral bending, as we expected an increase on the ipsilateral LD due to its lateral flexion function. This could suggest that static lateral bending of the trunk is generated mainly due to RA contraction. However, this needs synchronised kinematics and sEMG studies to be able to reach a conclusive result. The contralateral LD contracting during lateral bending can be explained by its phasic activation, and the fact that LD seems to contract in response to back destabilisation as seen in trot [25]. Activation in trot is higher during the stance phase of the contralateral forelimb than for the ipsilateral one due to the relative instability of the trunk being counteracted by activity in the trunk muscles [25]. The lower level of peak and ARV sEMG on LD during spinal flexion exercises was due to eccentric contraction in order to prevent hyper flexion. This data regarding LD activation differs from data collected by pushing a blunt object on the LD to elicit lateral flexion[26] showing that reflex activation differs from voluntary activation, even when the same movement is sought. The lack of significant findings in the LD in the lateral bending exercises suggest that other muscles actively induce latero-flexion of thoracolumbar region during DME rather than what was found on induced reflex lateral bending [18,26].

A trend to increase LS flexion gradually from chin-to-chest followed by chin-to-carpus and chin-to-fetlock was observed, although differences were not significant. In agreement with previous studies [23]no differences were found on the lumbar spine motion during different flexion exercises, although the greatest differences were seen on the cervical vertebrae and the thoracic spine, with chin-to-carpi having the greatest flexion on the cranial thoracic vertebrae (T6 to T8) while chin-to-fetlock had a greater angulation of the caudal thoracic (T10-T16).

Pelvic lift anecdotally stimulates the abdominal and sublumbar muscles to flex and lift the lumbar and LS joints [2]. Our results confirm that pelvic lift generated significantly greater lumbo-sacral flexion compared to thoracic lift (p= 0.031). Conversely, RA muscle activation was found to be greater during thoracic lift exercises. The same findings were observed on the peak muscle contraction in other studies [17]. It could be hypothesized that RA muscle is activated when performing exercises involving a greater flexion of the thoracic and cranial lumbar vertebrae, and, based on equine anatomy the *iliopsoas* complex would play a larger role in lumbo-sacral flexion, validating the importance of this exercise. In agreement with Barsanti et al. [17] a greater LD muscle activation was seen on the thoracic lift compared to the pelvic lift, although not significant. Like in the other exercises, LD activation could be associated to the eccentric contraction to accomplish its antagonist function or concentric contraction to return to neutral position. Overall, pelvic lift showed the greatest lumbo-sacral flexion, yet RA and LD EMG amplitude signal were lower than thoracic lift, suggesting that sub-lumbar muscles have a greater contribution to flex lumbo-sacral region than the muscles assessed in this study.

Previous studies have focused on the long-term benefit of DME in increasing and improving CSA of *m. multifidus* [6–8]. The current study found an increase of RA muscle activity, which could contribute to RA hypertrophy as reported by Rodrigues et al. [6]. This finding contributes to support DMEs for core strengthening but further studies should evaluate the effect of a long term DME on another core musculature.

RA is a highly important back stabilizing muscle[1]. In relation to the bow and string concept, RA strengthening could potentially improve back flexion and consequently hind limb protraction. Strengthening of this muscle could also help to reduce back pain and promote long career longevity in equine athletes. Results on RA, in this study, were consistent, giving a guideline of the level of muscle effort required in each exercise. This could enable physiotherapists and veterinarians to prescribe more efficient and less time-consuming rehabilitation plans.

Nine horses were included in our research, in line with existing studies. Exercises chosen were based on Stubbs and Clayton [2] and exercises were performed following the book’s guidelines with some adaptations, as. spinal bending exercises differ from the ones described, in the intensities. This decision was made in consideration of each horses’ welfare, as chin-to-fetlock lateral bending exercise require a certain level of fitness and training in order to perform the exercises correctly.

This study brings a contribution in elucidating how one epaxial, one hypaxial muscle and the lumbosacral joint responds to different core strengthening exercises. We have identified that different exercises have different benefits, so a core strengthening programme including a variety of exercises would be recommended. Due to the clear increased muscle activity with the progression of reach between exercises, we would recommend an increase in exercise level when the horse seems comfortable performing the previous level.

The main limitation of this study is that we have measured sEMG only in one side, without accounting for laterality. We have tried to mitigate this issue by measuring the muscle on lateral bending to both sides, but on an ideal scenario, measurements should have been taken from both sides simultaneously to understand effects of laterality. Furthermore, for the belly lifts, as the pression made by the hands is relatively close to the sEMG electrodes, we cannot discard that some of the measurements had motion artifacts. Finally, we have observed an immense inter-horse variation on the amount of muscle activity whilst performing the exercises, which could be due to different core strength and fitness amongst horses. However, this study design emphasised comparisons within each horse between exercises. A question that remains is how muscle activation would change following a long-term core strengthening training.

In conclusion, the RA has been proved to be highly targeted with spinal flexions, lateral bending and thoracic lifts. Pelvic lift exercises are beneficial in flexing the lumbo-sacral joint, which is paramount for joint health and horse performance. Core strengthening exercises should be recommended as routine for core strengthening, helping in injury prevention, as well as part of rehabilitation protocols.

